# Human club cells derived from pluripotent stem cells reveal new insights into epithelial lineage plasticity through structural and functional validation

**DOI:** 10.64898/2026.05.20.726364

**Authors:** Naoyuki Sone, Naoki Fujiwara, Abeer Keshta, Satoshi Konishi, Manabu Toyoshima, Tomoyuki Takaku, Yasuhiko Takahashi, Mio Iwasaki, Takuya Yamamoto, Shimpei Gotoh

**Affiliations:** Center for iPS Cell Research and Application (CiRA), Kyoto University, Kyoto 606-8507, Japan; Environmental Health Science Laboratory, Sumitomo Chemical Co., Ltd., Osaka 554-8558, Japan; Institute for the Advanced Study of Human Biology (WPI-ASHBi), Kyoto University, Kyoto 606-8501, Japan; Medical-risk Avoidance Based on iPS Cells Team, RIKEN Center for Advanced Intelligence Project (AIP), Kyoto 606-8507, Japan

## Abstract

Airway epithelial homeostasis relies on multiple specialized cell types, with club cells playing central roles in maintaining epithelial integrity and regulating inflammation. Environmental insults such as allergens, viral infections, or pollutants preferentially damage club cells, impairing epithelial repair and contributing to pulmonary diseases. However, the functional properties of club cells remain incompletely defined, and tractable human models are lacking. Herein, we establish a robust platform to differentiate human pluripotent stem cells (hPSCs) into club cells exhibiting their hallmark secretory features, appropriate epithelial organization, and functional properties. Single-cell transcriptomic analyses and lineage trajectory inference revealed unexpected epithelial plasticity: hPSC-derived club cells give rise to multiciliated epithelial cells through a deuterosomal intermediate—a previously uncharacterized trajectory. Additionally, a distinct club cell subset exhibited transcriptional features indicative of neuroendocrine and goblet cell differentiation potential. This study uncovers club cell plasticity and establishes a hPSC-based platform for studying airway development, regeneration and disease modeling.

---

The human airway epithelium preserves tissue integrity and regenerative capacity through coordinated interactions among various epithelial cell states. Among these, club cells constitute a population of non-ciliated secretory epithelial cells predominantly located in the distal airways that contribute to airway homeostasis by facilitating mucociliary clearance through the secretion of airway lining fluid constituents ^1^ and immunomodulatory factors. Beyond their essential roles in epithelial maintenance, club cells are critically involved in xenobiotic metabolism and detoxification via the cytochrome P450 system, and they modulate immune and inflammatory responses by releasing club cell secretory protein (CCSP/SCGB1A1) and other mediators ^2-5^. Additionally, club cells exhibit progenitor-like properties, including self-renewal and the capacity to differentiate into multiciliated cells during tissue homeostasis and repair ^3,6,7^.

Murine studies have demonstrated that extensive basal cell depletion promotes the dedifferentiation of club cells into basal-like progenitors, which subsequently expand to facilitate epithelial regeneration ^8^. Additionally, bronchioalveolar stem cells (BASCs), defined by co-expression of SCGB1A1 and SFTPC, contribute to the regeneration of bronchiolar and alveolar epithelial lineages at the bronchioalveolar duct junction ^9-11^. Recent single-cell transcriptomic analyses have identified SCGB3A2-positive secretory cell populations enriched in the terminal respiratory bronchioles, underscoring regional and molecular heterogeneity within the airway secretory compartment ^12,13^. However, the mechanisms governing the establishment, maintenance, and interconversion of these distinct epithelial states in the human airway remain incompletely understood, primarily due to the absence of robust, physiologically relevant human model systems.

Despite their critical physiological roles, club cells are highly susceptible to environmental and chemical insults. Exposure to cigarette smoke ^14^, photochemical smog ^15^, toxic chemicals ^16,17^, and pharmacological agents, including anticancer therapies ^18^, preferentially compromise club cells, impairing epithelial repair and barrier function. Accordingly, *CCSP* serves as a widely recognized biomarker for airway and alveolar epithelial injury ^19,20^. Club cell injury has been implicated in the pathogenesis of chronic obstructive pulmonary disease (COPD) ^21^ and bronchiolitis obliterans following lung transplantation ^22^, in which loss or dysfunction of club cells is associated with sustained inflammation and impaired epithelial regeneration. However, there remains a paucity of experimental systems that accurately recapitulate the secretory and regenerative functions of human club cells, thus limiting investigations into their contributions to human airway diseases ^23^.

Given the central role of club cells in airway biology and pathology, robust experimental models are required to elucidate their functional properties and responses to injury. While club-like cells have been successfully differentiated from primary human airway epithelial cultures ^24-26^ and from human PSCs (hPSCs) ^12,27-31^, existing systems are constrained by variability, incomplete functional assessment, and limited scalability. Furthermore, current models often fail to capture the developmental plasticity and potential of human club cells, or to provide a tractable framework for disease modeling relevant to human physiology.

In this study, we sought to establish a robust, reproducible differentiation platform to generate structurally and functionally validated hPSC-derived club cells, supported by transcriptomic and protein-level validation. To this end, we generated an SCGB1A1 reporter hPSC line, a canonical marker of airway club cells, and used fluorescent protein–based tracking to optimize and validate the differentiation protocol. The resulting club cells were then comprehensively characterized using complementary analytical approaches to systematically assess their structural and functional properties and to explore additional aspects of club cell biology. Through this integrated strategy, hPSC-derived club cells are established as a robust and physiologically relevant model for investigating airway epithelial development, regeneration, disease mechanisms.

## Results

### Generation of SCGB1A1-positive club cells from hPSCs

Club cells are essential regulators of airway epithelial homeostasis, xenobiotic detoxification, and epithelial regeneration. However, progress in mechanistic and translational studies has been hampered by the absence of reliable, reproducible methods for generating or expanding human club cells from pluripotent stem cells or primary sources ^6,26,32,33^. To overcome this limitation, we aimed to develop a differentiation platform that enables efficient induction, purification, and quantitative analysis of *SCGB1A1*-expressing club cells derived from hPSCs.

For precise monitoring of club cell specification, we engineered an *SCGB1A1* reporter hPSC line via targeted knock-in of a green fluorescent protein (GFP) cassette at the endogenous locus (Fig. 1a; Extended Data Fig. 1a, b). Using this reporter hPSC line, we established a stepwise differentiation protocol incorporating stage-specific Notch signaling activation from day 21 onward (Fig. 1b), aiming to recapitulate Notch’s critical function in promoting secretory lineage commitment within the distal airway epithelium during development and regeneration ^34,35^. Consistent with previous studies demonstrating that Jagged-1-mediated Notch signaling is required for club cell differentiation from basal cells *in vivo* and in primary human airway epithelial cultures ^36,37^, we systematically optimized culture conditions, as well as the concentration and type of Notch ligands applied (Extended Data Fig. 1c, d).

**Fig. 1:**
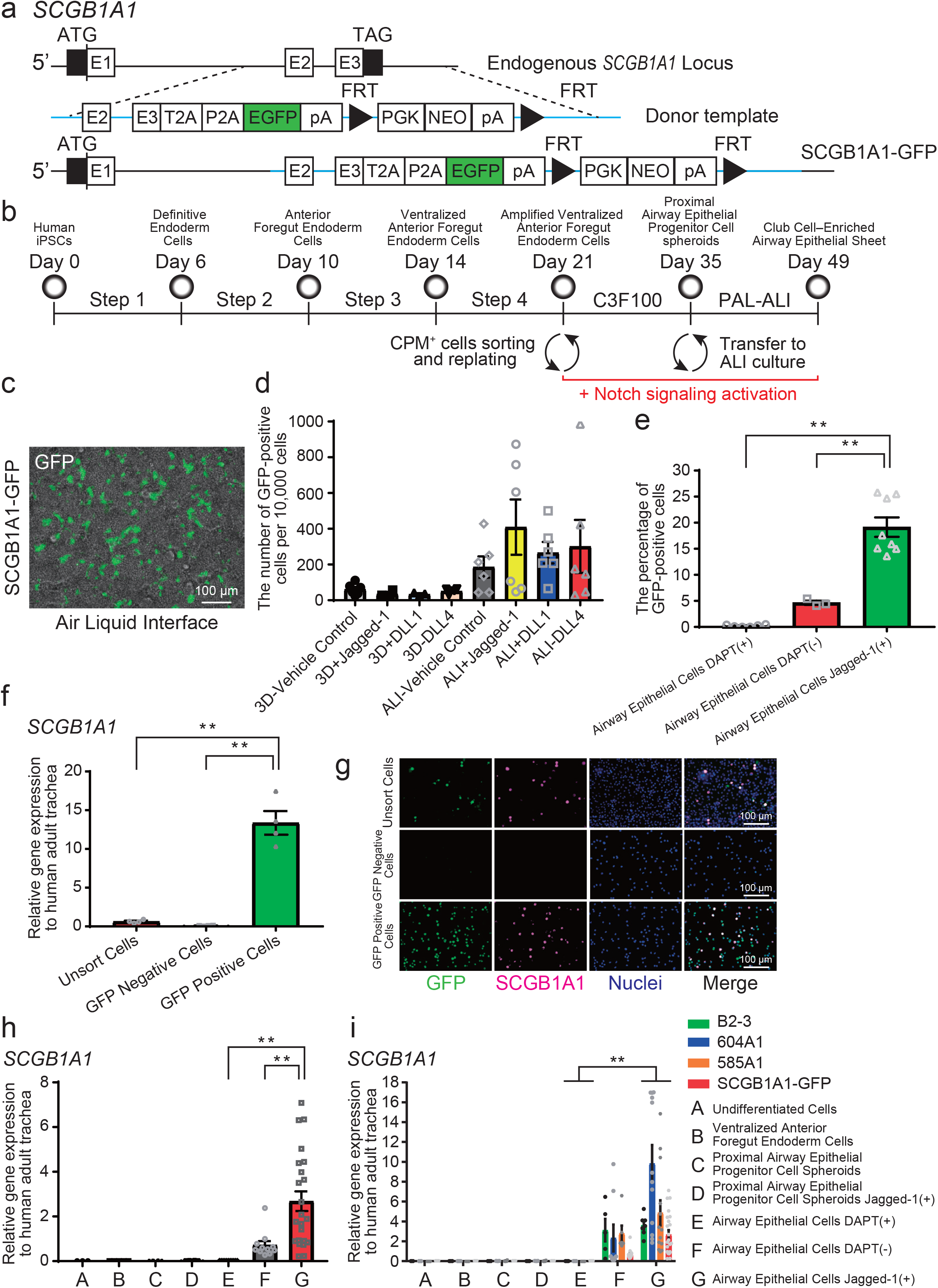
Establishment of a robust and reproducible differentiation protocol for SCGB1A1-positive club cells derived from hPSCs. (a) Schematic design of an SCGB1A1 reporter hPSC line. (b) Schematic overview of the club cell differentiation protocol with stage-specific Notch signaling activation initiated from day 21. (c) Validation of GFP expression during differentiation into club cells. (d) Comparative flow cytometric analysis of GFP expression during club cell differentiation in 3D organoid and ALI cultures (mean ± SEM, *n* ≥ 3 independent experiments). (e) Comparison of GFP-positive cell ratios among three club cell differentiation conditions (mean ± SEM, *n* ≥ 3 independent experiments; one-way ANOVA with Tukey’s multiple comparisons test; \*\**p* < 0.01). (f) RT-qPCR analysis of *SCGB1A1* expression in FACS-sorted GFP-positive and GFP-negative populations compared with unsorted cells (mean ± SEM, *n* ≥ 3 independent experiments; one-way ANOVA with Tukey’s multiple comparisons test; \*\**p* < 0.01). (g) Validation of SCGB1A1 protein expression in FACS-sorted GFP-positive and GFP-negative cells by immunofluorescence staining. (h) RT-qPCR evaluation of each club cell differentiation protocol within the SCGB1A1 reporter cell line (mean ± SEM, *n* ≥ 3 independent experiments; one-way ANOVA with Tukey’s multiple comparisons test; \*\**p* < 0.01). (i) RT-qPCR evaluation of the robustness and reproducibility of the club cell differentiation protocol across multiple normal hPSC lines (mean ± SEM, *n* ≥ 3 independent experiments; two-way ANOVA with Dunnett’s multiple comparisons test; \*\**p* < 0.01).

The final protocol involved the addition of Jagged-1 at 10 ng/mL under air–liquid interface (ALI) conditions. GFP expression increased progressively over the course of cell differentiation (Fig. 1c, Extended Data Fig. 1c), serving as a quantitative indicator to optimize differentiation efficiency. Flow cytometric analysis revealed that *SCGB1A1*-positive cells were more effectively induced in ALI cultures compared to three-dimensional (3D) organoid systems, with Notch activation further augmenting the proportion of GFP-positive cells (Fig. 1d, Extended Data Fig. 1d). The revised protocol significantly improved induction efficiency of club cells relative to conventional methods (Fig. 1e), while also supporting prolonged cell maintenance (Fig. S1e).

Fluorescence-activated cell sorting demonstrated significant enrichment of SCGB1A1 transcripts in GFP-positive cells relative to GFP-negative and unsorted populations (Fig. 1f), confirming the specificity of the reporter system. Immunofluorescence assays corroborated robust SCGB1A1 protein expression exclusively within GFP-positive cells (Fig. 1g, Extended Data Fig. 1f), while quantitative RT–PCR confirmed consistent induction of SCGB1A1 and other club cell markers following the established protocol (Fig. 1h), illustrating the scalability of this platform. Notably, the optimized differentiation protocol was robust and reproducible across multiple independent hPSC lines (Fig. 1i, Extended Data Fig. 1g).

Collectively, these results establish a robust and reproducible framework for generating SCGB1A1-positive club cells from hPSCs, providing a foundation for subsequent functional validation, transcriptomic profiling, and lineage trajectory analyses.

### hPSC-derived club cells recapitulate the structural and functional hallmarks of airway club epithelium

To evaluate the maturation of hPSC-derived club cells produced via our established protocol, we conducted a detailed phenotypic, ultrastructural, and transcriptomic analysis of club cell-enriched airway epithelial sheets. Immunofluorescence analysis demonstrated that SCGB1A1-positive cells were distributed in a mosaic-like pattern throughout the epithelial sheet (Fig. 2a, Extended Data Fig. 2a), closely resembling the spatial arrangement of club cells in bronchial tissue *in vivo* ^34,38^. Quantitative immunostaining confirmed that the proportion and distribution of scattered *SCGB1A1*-positive club cells within these sheets closely resembled those in bronchial tissue (Fig. 2b, Extended Data Fig. 2b) ^5,39^.

**Fig. 2:**
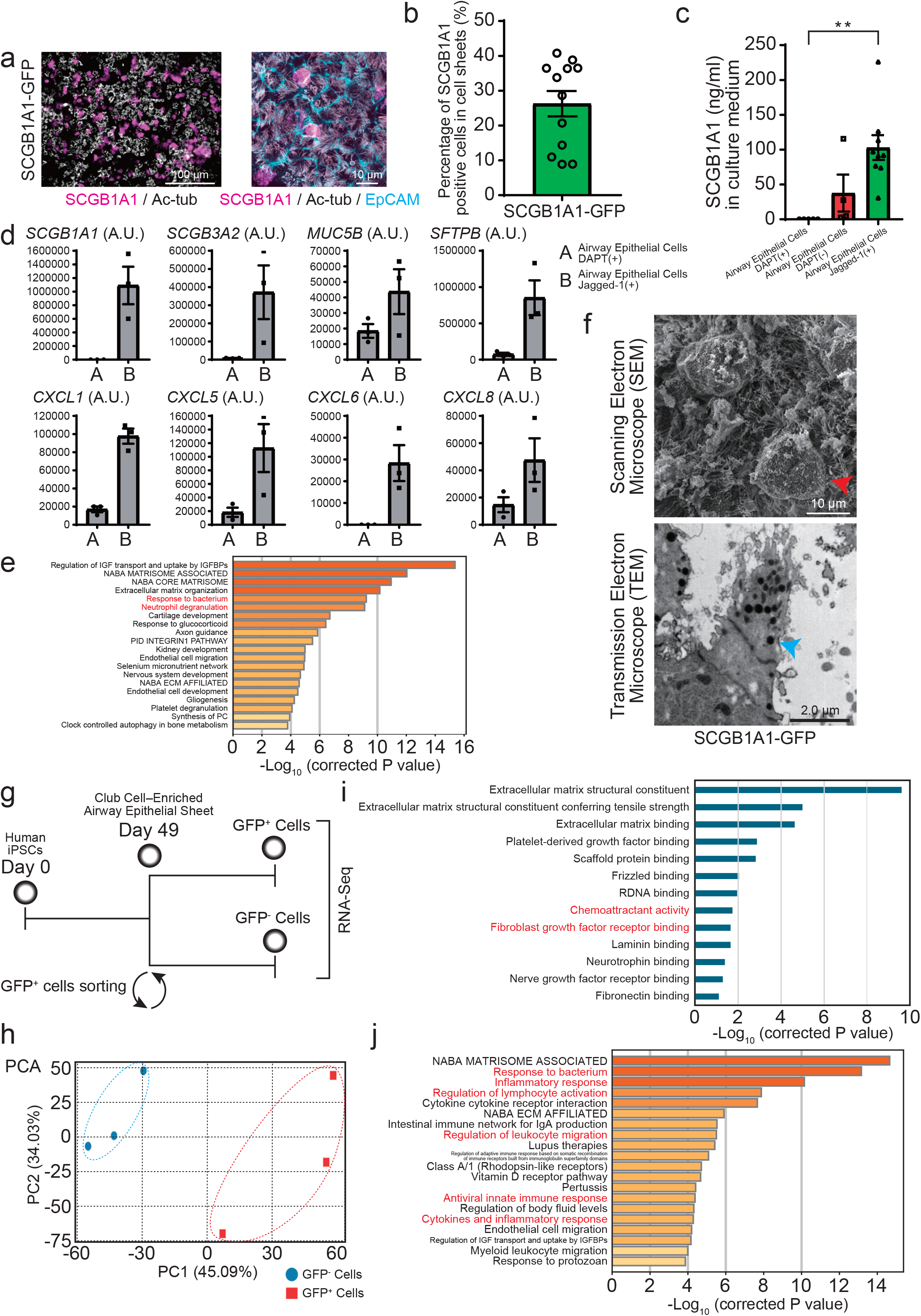
Structural and functional maturation of the club cell-enriched airway epithelial sheet generated using the established protocol. (a) Scattered distribution of SCGB1A1-positive club cells within the club cell-enriched airway epithelial sheet. (b) Quantification of the proportion of scattered SCGB1A1-positive club cells in the airway epithelial sheet by immunofluorescence staining (mean ± SEM, *n* ≥ 3 independent experiments). (c) Secretory function of airway epithelial sheets with and without Jagged-1 supplementation using SCGB1A1 ELISA, compared to the effect of Notch inhibitor, DAPT supplementation (mean ± SEM, *n* ≥ 3 independent experiments; one-way ANOVA with Tukey’s multiple comparisons test; \*\**p* < 0.01). (d) Comparative secretome analysis of the club cell-enriched airway epithelial sheet and a conventional airway epithelial sheet (mean ± SEM, *n* = 3 independent experiments). Values are presented in protein area (A.U.), derived from mass spectrometry-based proteomics. (e) Metascape-based enrichment analysis of secretome differences between the club cell-enriched airway epithelial sheet and a conventional airway epithelial sheet. (f) Ultrastructural characterization of the club cell-enriched airway epithelial sheet by transmission and scanning electron microscopy; red arrows: club cells, blue arrows: secretory granules. (g) Schematic overview of RNA-seq analysis of GFP-positive and GFP-negative populations isolated from the club cell-enriched airway epithelial sheet. (h) Principal component analysis of RNA-seq data from GFP-positive and GFP-negative populations. Gene Ontology enrichment analysis of RNA-seq data from GFP-positive and GFP-negative populations. Metascape-based enrichment analysis of RNA-seq data from GFP-positive and GFP-negative populations.

Secretory function was assessed via enzyme-linked immunosorbent assays (ELISAs), revealing robust secretion of SCGB1A1 from the club cell-enriched airway epithelial sheets, with significantly higher levels than the Notch-suppressed hPSC-derived airway epithelial sheets lacking directed club cell induction (Fig. 2c, Extended Data Fig. 2c). Unbiased secretome analysis further demonstrated secretion of club cell-associated proteins SCGB1A1 and SCGB3A2, as well as MUC5B, SFTPB, and various chemokines (Fig. 2d, Extended Data Fig. 2D–F). Functional annotation using Metascape highlighted enrichment of biological processes characteristic of club cells, including regulation of inflammatory responses (Fig. 2e).

Transmission (TEM) and scanning electron microscopy (SEM) revealed that cells within the club cell-enriched airway epithelial sheet exhibited dome-shaped apical morphology and abundant electron-dense secretory granules, features indicative of mature club cells (Fig. 2f, Extended Data Fig. 2g). Transcriptomic profiling was performed by isolating GFP-positive and GFP-negative populations from club cell-enriched airway epithelial sheets derived from SCGB1A1 reporter hPSCs, followed by bulk RNA sequencing (Fig. 2g). Principal component analysis (PCA) demonstrated clear segregation between GFP-positive and GFP-negative populations (Fig. 2h), reflecting distinct transcriptional profiles. GFP-positive cells were enriched for gene markers characteristic of club cells (Extended Data Fig. 3a–c). Further Gene Ontology and Metascape analyses revealed that GFP-positive cells were selectively enriched for gene programs associated with secretory activity, regulation of inflammatory responses, and airway-specific defense mechanisms (Fig. 2i, j). These structural, functional, and transcriptomic features were reproduced across multiple independent hPSC lines (Extended Data Fig. 2a–c, g), confirming the protocol’s robustness and reproducibility across genetic backgrounds.

Collectively, these results demonstrate that hPSC-derived club cells generated through our protocol accurately recapitulate the key architectural organization, secretory functions, ultrastructural features, and molecular identity of native airway club epithelium.

### Single-cell trajectory analysis identifies two differentiation routes from club cells: multiciliated and neuroendocrine lineages

To clarify the molecular mechanisms driving the lineage plasticity of hPSC-derived club cells, we performed single-cell RNA sequencing (scRNA-seq) of the club cell-enriched airway epithelial sheet (Fig. 3a). Transcriptomic profiling identified several distinct epithelial cell clusters within the differentiated sheet, including SCGB1A1^+^ Proliferating club cells, terminal respiratory bronchiole secretory cell (TRB-SC)-like cells, SCGB1A1^+^ club cell, deuterosomal cells ^40^, multiciliated epithelial cells, and neuroendocrine cells (Fig. 3b).

**Fig. 3:**
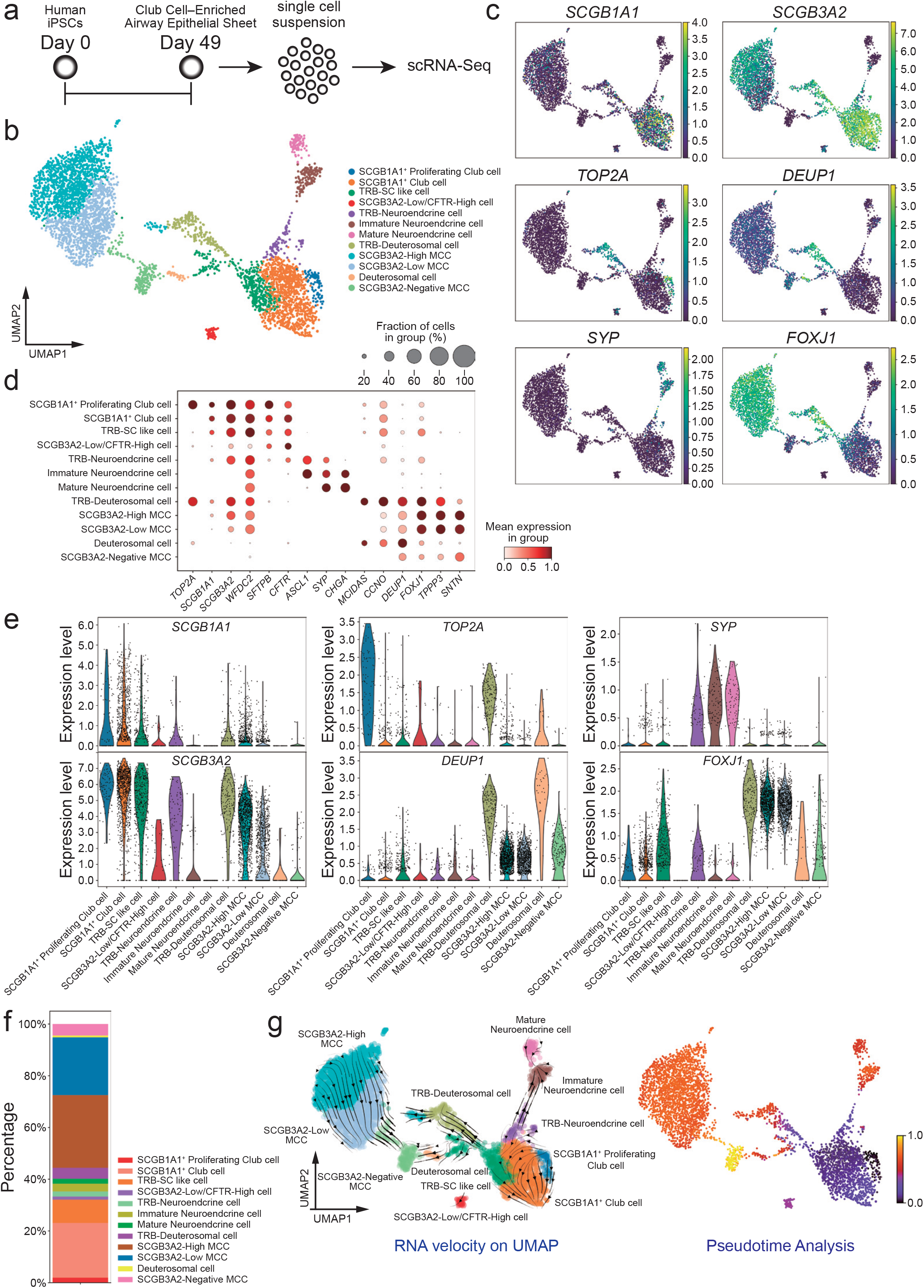
Single-cell transcriptomic discovery of SCGB3A2-positive club cell differentiation trajectories toward neuroendocrine and multiciliated epithelial lineages via deuterosomal intermediates. (a) Schematic overview of the scRNA-seq workflow following differentiation into the club cell-enriched airway epithelial sheet. (b) Clustering analysis of single-cell transcriptomic data derived from the club cell-enriched airway epithelial sheet. (c) Feature plot analysis of representative marker genes identifying club cells, SCGB1A1+ proliferative cells, deuterosomal cells, multiciliated epithelial cells, and neuroendocrine cells. (d) Dot plot analysis of representative marker gene expression across clusters identified in the club cell-enriched airway epithelial sheet. (e) Violin plot analysis of representative marker gene expression across cell clusters identified in the club cell-enriched airway epithelial sheet. (f) Bar plot showing the relative proportions of cell clusters identified in the club cell-enriched airway epithelial sheet. (g) Pseudotime trajectory analysis of scRNA-seq data mapped onto UMAP space of the identified cell clusters, color-coded by pseudotime values.

Robust cluster annotation was achieved through feature plot and dot plot analyses using established lineage markers (Fig. 3c, d, Extended Data Fig. 4a). Club cell clusters expressed canonical secretory markers, including *SCGB1A1* and *SCGB3A2*, whereas proliferative cells expressed cell cycle-associated genes such as *TOP2A*. A discrete deuterosomal population was defined by enrichment of *DEUP1, CCNO*, and *MCIDAS—*genes indicative of intermediate stages in multiciliated cell differentiation ^41^. Fully differentiated multiciliated epithelial cells highly expressed *FOXJ1* and *TPPP3*. Additionally, a distinct cluster expressing neuroendocrine-associated genes, including *SYP, ASCL1*, and *CHGA*, was identified, indicating the emergence of a neuroendocrine lineage within the club cell-enriched sheet (Fig. 3c–e, Extended Data Fig. 4a, b).

Quantitative analysis of cluster proportions in our dataset demonstrated that club-related cells constituted the substantial population within the enriched airway epithelial sheet (Fig. 3f), closely recapitulating the relative abundance reported for club cells in human bronchial epithelium in vivo ^39^. Additional populations, including deuterosomal cells, multiciliated epithelial cells, and neuroendocrine cells appeared at appropriate relative proportions, confirming that the differentiated sheet recapitulated the cellular diversity of native airway epithelium, encompassing major and rare epithelial lineages (Fig. 3f) ^13,42^. Notably, *SCGB3A2* expression was enriched in a subset of club cells positioned transcriptionally between the canonical club cell cluster and downstream differentiated lineages, suggesting the existence of a transitional progenitor-like population.

The single-cell transcriptomic analysis yielded two key observations. First, two distinct deuterosomal cell clusters were identified: one expressing *SCGB3A2* and another lacking its expression. Similarly, multiciliated epithelial cells (MCCs) were categorized into SCGB3A2-expressing and SCGB3A2-non-expressing subpopulations, representing transitional states downstream of SCGB3A2-positive deuterosomal cells. SCGB3A2-high MCCs were distributed across multiple clusters displaying graded expression levels. Although SCGB3A2-high MCCs appeared as three spatially separated groups in UMAP space, they exhibited a shared transcriptional signature and thus were classified collectively as a single SCGB3A2-high MCC cluster.

Pseudotime trajectory analysis supported a sequential differentiation continuum originating from club cells, progressing through TRB-like secretory intermediates and SCGB3A2-positive deuterosomal cells to SCGB3A2-high MCCs, with a subsequent gradual transition toward SCGB3A2-low and ultimately SCGB3A2-negative MCCs (Fig. 3g). Additionally, an alternative trajectory was observed wherein SCGB3A2-negative deuterosomal cells transitioned directly to SCGB3A2-negative MCCs, suggesting the presence of multiple routes for MCC differentiation. These results support a stepwise multiciliogenesis program in which *SCGB3A2* expression is maintained during the early stages of MCC differentiation and progressively downregulated during terminal maturation.

Second, we identified a distinct cluster of *SCGB3A2*-positive neuroendocrine cells. Pseudotime trajectory analysis demonstrated that this lineage arose from club cells, passing through an *SCGB3A2*-positive neuroendocrine intermediate state before differentiating into mature neuroendocrine cells (Fig. 3g). This trajectory suggests a previously unappreciated differentiation route linking club cells to the neuroendocrine lineage via a transient TRB-neuroendocrine cell intermediate.

Collectively, these single-cell transcriptomic analyses reveal the lineage plasticity of hPSC-derived club cells and identify *SCGB3A2*-positive club cells as a key transitional population linking secretory club cells to multiciliated and neuroendocrine epithelial lineages.

### hPSC-derived club cells exhibit multilineage differentiation potential toward multiciliated, neuroendocrine, and alveolar epithelial fates

To validate the multilineage differentiation potential inferred from single-cell transcriptomic analyses, we conducted immunofluorescence-based protein characterization of hPSC-derived club cells and their differentiated progeny.

Consistent with the transcriptomic heterogeneity, we identified SCGB1A1^+^/SCGB3A2^+^/MUC5B^+^ secretory cells, SCGB1A1^+^/SFTPB^+^ cells with alveolar-associated features, and SCGB1A1^+^/SCGB3A2^+^/Ki-67^+^ proliferative club cells. This demonstrates the coexistence of secretory, alveolar-like, and proliferative club cell subpopulations at the protein level (Fig. 4a). Immunofluorescence staining confirmed TRB-associated deuterosomal cells co-expressing SCGB1A1, FOXJ1, and CCNO, as well as TRB-neuroendocrine cells expressing SCGB1A1, SCGB3A2, and SYP (Fig. 4b). Additionally, SCGB3A2-high MCCs were characterized by acetylated tubulin and FOXJ1, providing protein-level evidence for differentiation toward the multiciliated lineage via a deuterosomal intermediate state (Fig. 4b).

**Fig. 4:**
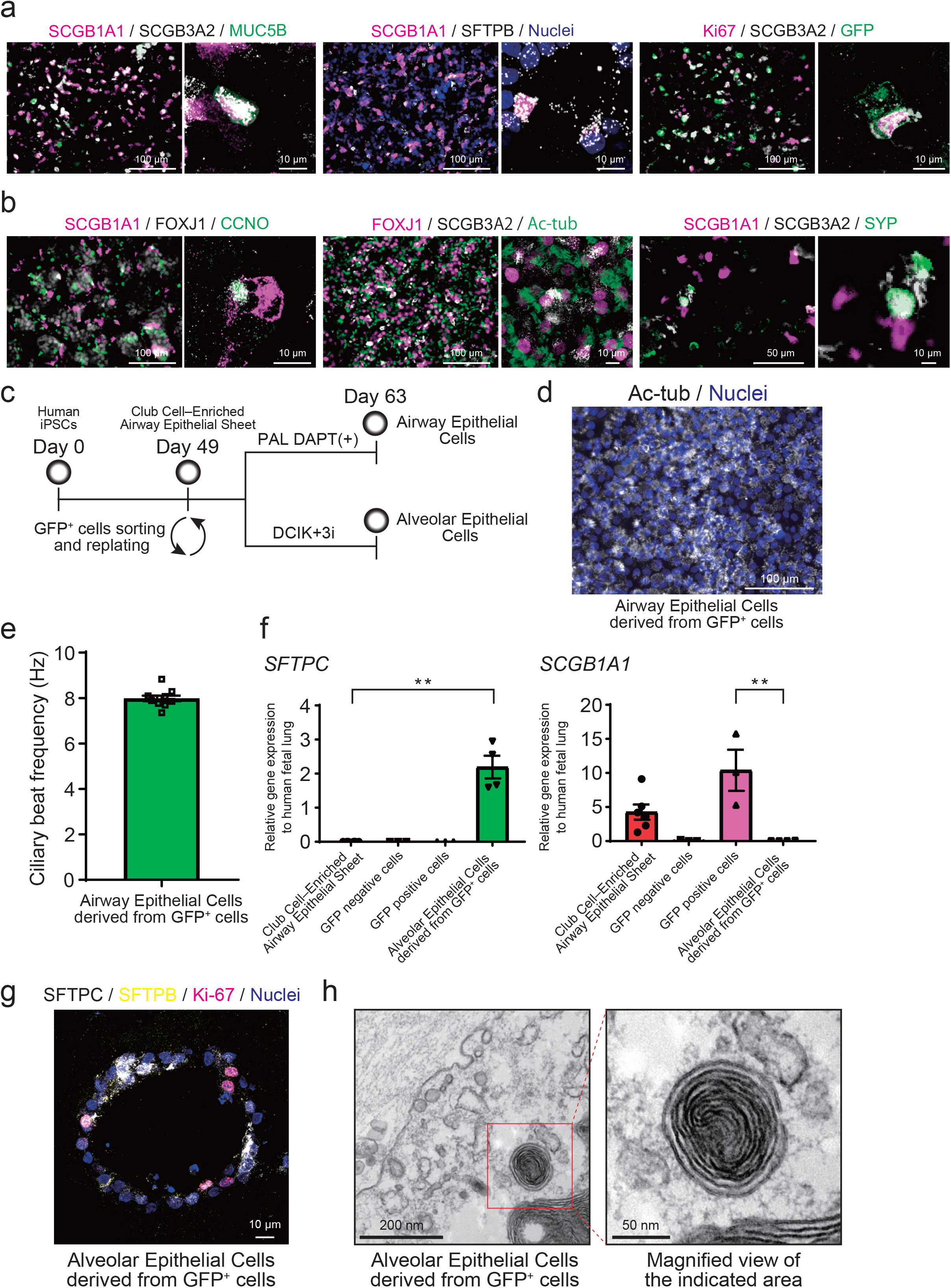
Extended lineage plasticity of hPSC-derived club cells toward multiciliated, neuroendocrine, and alveolar epithelial lineages. (a) Immunofluorescence validation of diverse hPSC-derived club cell subpopulations. (b) Immunofluorescence validation of TRB-deuterosomal cells, TRB-neuroendocrine cells, and SCGB3A2-high multiciliated epithelial cells identified by single-cell transcriptomic analysis. (c) Schematic overview of the isolation of GFP-positive club cells and their subsequent differentiation toward multiciliated epithelial cells and alveolar type 2 cells. (d) Immunofluorescence validation of multiciliated epithelial cell differentiation from GFP-positive club cells. (e) High-speed video microscopy analysis of ciliary beat frequency in multiciliated epithelial cells differentiated from GFP-positive club cells. (f) RT-qPCR analysis of alveolar type 2 cell differentiation from GFP-positive club cells, including SCGB1A1 expression, following differentiation (mean ± SEM, *n* ≥ 2 independent experiments; one-way ANOVA with Tukey’s multiple comparisons test; \*\**p* < 0.01). (g) Immunofluorescence validation of alveolar type 2 cell differentiation from GFP-positive club cells. (h) Electron microscopic identification of lamellar bodies in alveolar type 2 cells differentiated from GFP-positive club cells.

To determine whether hPSC-derived club cells possess stem cell-like properties beyond secretory function, we evaluated their capacity to diversify into airway and alveolar epithelial fates. Although airway club cells function as facultative progenitors that regenerate MCCs after injury and, in specific contexts, contribute to alveolar epithelial lineages ^8-11^, the bipotency of hPSC-derived club cells has remained unclear. To address this, we isolated GFP-positive club cells from the club cell-enriched airway epithelial sheet using fluorescence-activated cell sorting and subjected them to lineage-specific differentiation protocols toward MCCs or alveolar type II (AT2) cells (Fig. 4c, Extended Data Fig. 4c).

Under multiciliogenesis-inducing conditions, GFP-positive club cells robustly differentiated into cells expressing canonical MCC markers, including acetylated α-tubulin, as confirmed by immunofluorescence (Fig. 4d). High-speed video microscopy demonstrated these cells exhibited coordinated ciliary activity with frequencies comparable to functional human airway multiciliated epithelium (Fig. 4e, Extended Data Movie 1). Under AT2 differentiation conditions, RT-qPCR analysis revealed significant upregulation of key AT2-associated genes, including *SFTPC*, compared with the original club cell-enriched airway epithelial sheet (Fig. 4f). During AT2 differentiation, *SCGB1A1* expression progressively decreased, indicating a transition away from the club cell phenotype, while AT2-associated gene expression concomitantly increased (Fig. 4f). Immunofluorescence confirmed expression of SFTPC and SFTPB, supporting successful AT2 differentiation from hPSC-derived club cells (Fig. 4g). Furthermore, TEM revealed lamellar bodies—definitive ultrastructural features of AT2 cells—within the spheroids, providing morphological confirmation of AT2 identity (Fig. 4h).

Collectively, the findings of the current study demonstrate that hPSC-derived club cells exhibit remarkable lineage plasticity, spanning proximal airway and distal alveolar epithelial fates. This intrinsic plasticity positions hPSC-derived club cells as a physiologically relevant epithelial progenitor population and establishes a robust human *in vitro* platform for analyzing airway–alveolar transitions during development, regeneration, and disease.

### IL-13 induces goblet cell differentiation and epithelial remodeling in airway epithelial sheets

We next examined whether club cell–enriched airway epithelial sheets recapitulate asthma-associated epithelial remodeling. Interleukin-13 (IL-13), a central type 2 cytokine implicated in asthma pathogenesis ^43,44^, was applied to the epithelial sheets (Fig. 5a). RT–qPCR analysis revealed that NKX2.1 expression was maintained following IL-13 stimulation, whereas expression of the club cell marker SCGB1A1 was reduced, accompanied by increased expression of goblet cell–associated genes, including MUC5AC and SPDEF (Fig. 5b, Extended Data Fig. 5a). These findings indicate a shift from club cell identity toward a goblet cell–like phenotype, consistent with epithelial remodeling. Immunofluorescence analysis confirmed increased MUC5AC protein expression together with reduced SCGB1A1 expression following IL-13 stimulation (Fig. 5c). MUC5AC secretion tended to increase following IL-13 stimulation, as determined by ELISA (Fig. 5d, Extended Data Fig. 5b). These transcriptional and secretory responses associated with goblet cells were consistently observed across independent hPSC lines, supporting the reproducibility of the model (Extended Data Fig. 5a, b).

**Fig. 5:**
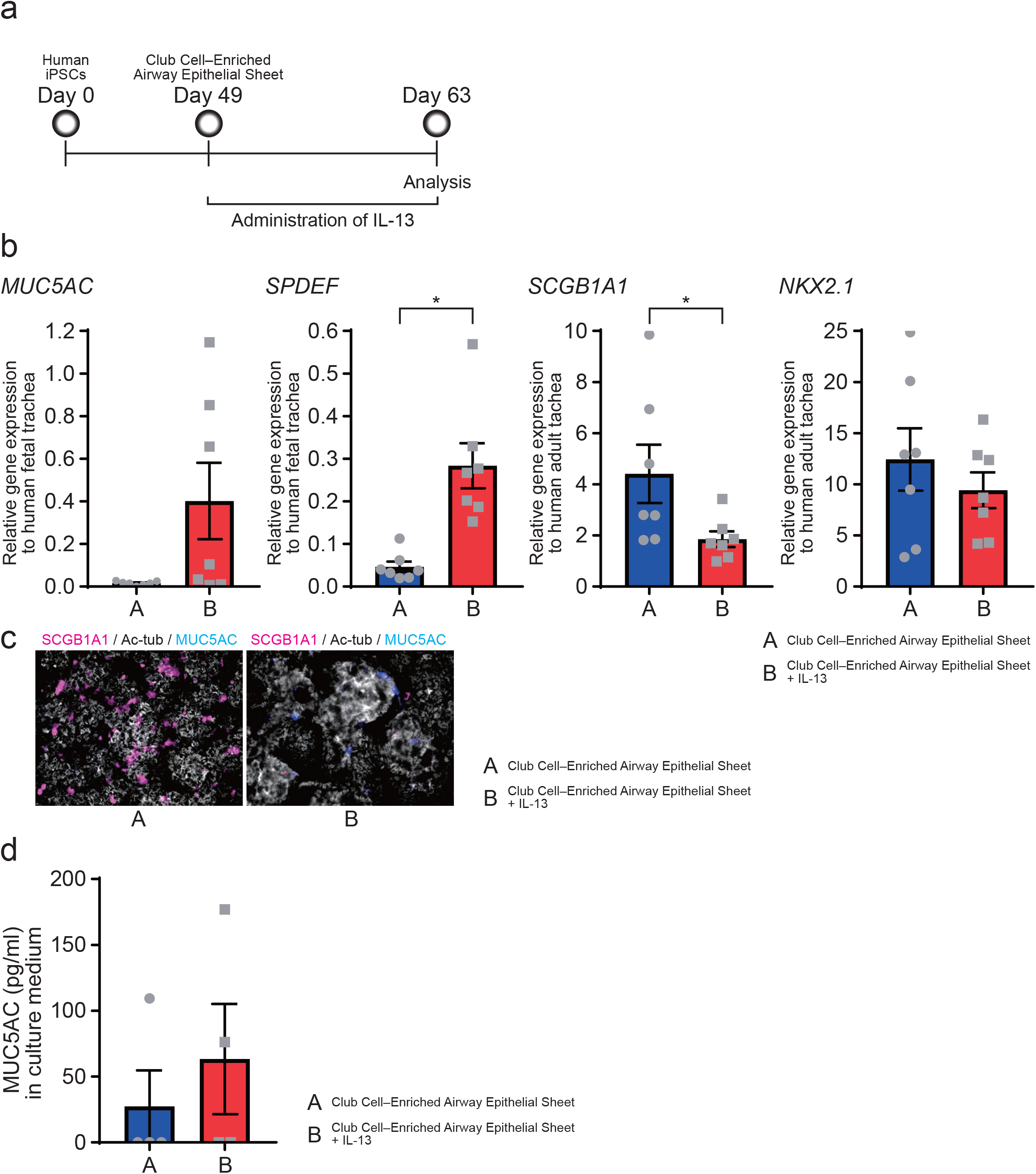
IL-13 induces goblet cell differentiation and epithelial remodeling in club cell–enriched airway epithelial sheets. (a) Schematic of IL-13 stimulation in club cell–enriched airway epithelial sheets. (b) RT–qPCR analysis of goblet and club cell marker gene expression in control and IL-13–treated airway epithelial sheets (mean ± SEM, n ≥ 3 independent experiments). (c) MUC5AC and SCGB1A1 protein expression in control and IL-13–treated airway epithelial sheets. (d) ELISA quantification of MUC5AC secretion in control and IL-13– treated airway epithelial sheets (mean ± SEM, n ≥ 2 independent experiments).

**Fig. 6:**
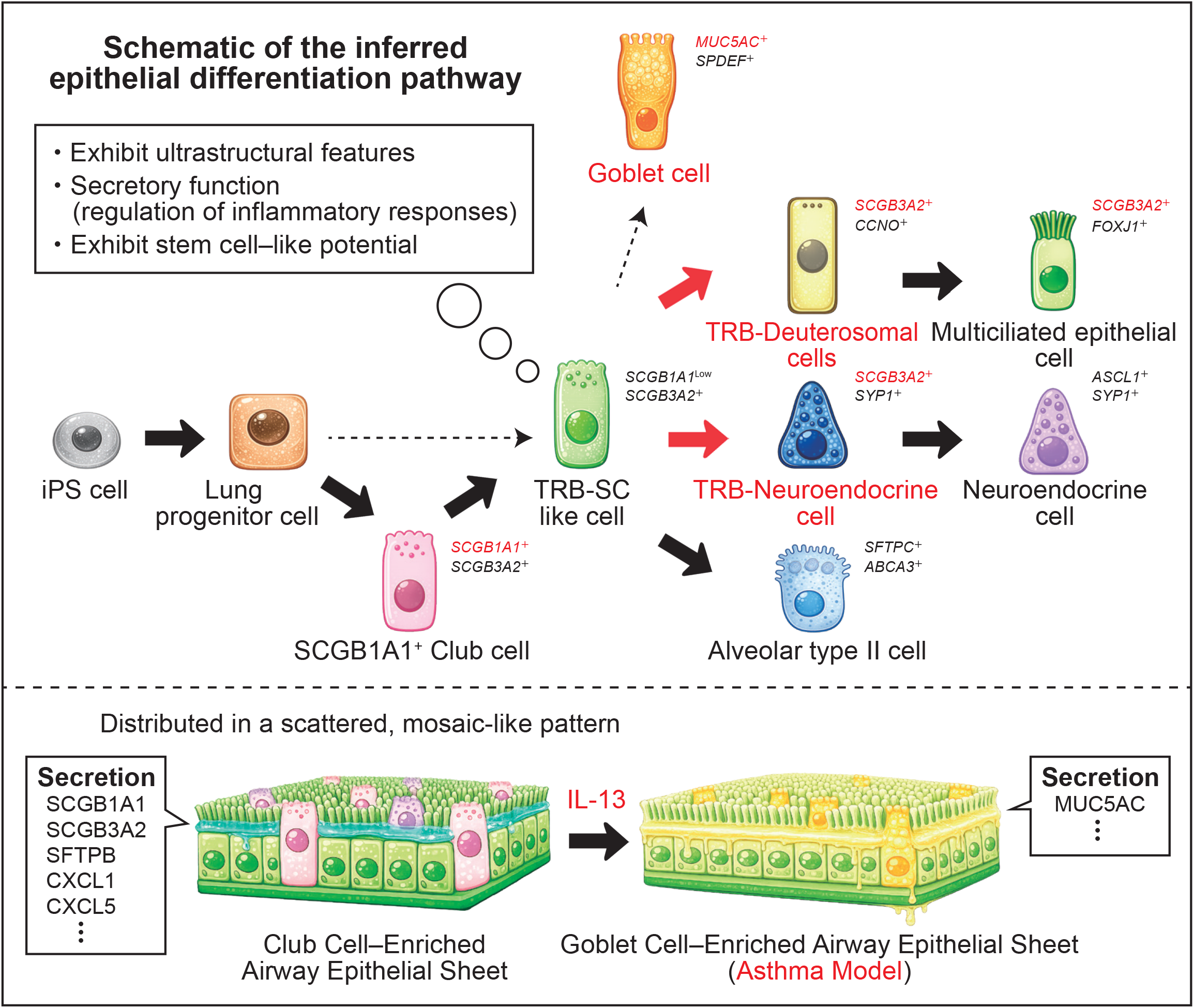
Summary of predicted epithelial differentiation pathways derived from hPSC-derived club cells. This schematic summarizes the predicted epithelial differentiation pathways identified in this study. hPSC-derived club cells exhibit structural and functional maturation and give rise to multiple airway epithelial lineages, including multiciliated epithelial cells via a deuterosomal intermediate, as well as a transcriptionally distinct subset with features suggestive of neuroendocrine differentiation potential. The diagram integrates findings from differentiation assays, single-cell transcriptomic analyses, and lineage trajectory inference to provide a conceptual overview of airway epithelial lineage relationships revealed by this study.

Collectively, these findings indicate that club cell–enriched airway epithelial sheets derived from hPSCs recapitulate key aspects of disease-associated epithelial remodeling, including IL-13–induced goblet cell differentiation. This platform thus provides a robust human *in vitro* model for investigating airway epithelial pathophysiology in a controlled and physiologically relevant context.

## Discussion

In this study, we developed a robust and reproducible platform for generating structurally and functionally validated airway club cells from hPSCs. By integrating reporter-based lineage tracing, quantitative functional assays, and single-cell transcriptomic profiling, our findings indicate that club cell-enriched airway epithelial sheets derived from hPSCs faithfully recapitulate the molecular characteristics, spatial organization, and functional properties of native human airway club epithelium. These results are consistent with documented epithelial lineage plasticity observed during injury-induced regeneration and disease-associated epithelial remodeling in the human lung ^45-49^.

Notably, single-cell analyses revealed previously underappreciated epithelial lineage plasticity, indicating that club cells can serve as progenitors for multiciliated and neuroendocrine lineages via distinct intermediate states. Furthermore, these engineered epithelial sheets respond to inflammatory stimuli such as IL-13 in a manner consistent with disease-associated airway remodeling, underscoring their potential for human-relevant disease modeling and therapeutic evaluation.

Our single-cell RNA sequencing and trajectory analyses provide evidence that human airway club cells can differentiate into MCCs through a *SCGB3A2*-positive deuterosomal intermediate. RNA velocity and pseudotime analyses consistently positioned *SCGB3A2*-expressing cells upstream of *FOXJ1*-positive MCCs, supporting a stepwise transition from club cell identity toward ciliogenic programs. This trajectory closely aligns with prior *in vivo* observations from single-cell analyses of the human distal airway epithelium ^13^, where transitional cell populations co-expressing club cell and ciliary markers were identified, and immunohistochemistry confirmed SCGB3A2 and *FOXJ1* co-expression *in situ*. The concordance between our hPSC-derived system and independent human airway datasets supports the physiological relevance of the club-to-ciliated differentiation axis reconstructed *in vitro*. Additionally, our results revealed a trajectory in which club cells transition through a TRB-positive neuroendocrine-like intermediate, suggesting a potential route toward pulmonary neuroendocrine cell specification. However, in contrast to the club-to-ciliated pathway, this lineage was not robustly validated using current human lung single-cell datasets. Publicly available scRNA-seq atlases of the human lung ^13,42^, including those derived from distal airways, consistently contain few neuroendocrine cells due to their inherent rarity and the technical challenges associated with their capture and preservation. Consequently, *SCGB3A2*-positive neuroendocrine populations are absent from these datasets, limiting direct *in vivo* corroboration of this differentiation route. Nevertheless, these findings underscore a potential advantage of club cell-enriched airway epithelial sheets derived from hPSCs: their capacity to model rare, transient, or developmentally restricted epithelial states that may be underrepresented or lost in analyses of primary tissues. Collectively, our data suggest that TRB-neuroendocrine intermediates derived from club cells likely constitute a difficult-to-observe lineage trajectory in the human airway epithelium.

Beyond developmental insights, our club cell-enriched airway epithelial sheets recapitulate hallmark features of disease-associated epithelial remodeling. These phenotypes parallel key pathological features observed in asthma, including mucus hypersecretion and epithelial remodeling driven by type 2 cytokines such as IL-13. Importantly, our data indicate that the platform can be further refined for therapeutic response modeling. For example, biologic agents targeting type 2 inflammatory pathways, such as IL-4 and IL-13 blockade with dupilumab ^50^, reduce mucus hypersecretion and improve airway function in patients with severe asthma. Incorporating such agents into this system would enable direct evaluation of epithelial-intrinsic therapeutic effects within a controlled human context, independent of systemic immune components. Consistent with this broader framework, previous research has established that airway club cells can generate alveolar epithelial lineages in parallel with ciliogenic differentiation, a process regulated by Notch-dependent mechanisms. The current study builds on these findings by using hPSC-derived club cells to validate known pathways and uncover novel aspects of lineage plasticity.

Limitations of this study warrant consideration. First, although our single-cell analyses strongly support club-to-ciliated and club-to-neuroendocrine differentiation trajectories, definitive *in vivo* lineage tracing in humans remains technically challenging. In particular, the scarcity of neuroendocrine cells in publicly available scRNA-seq datasets hinders robust validation of the proposed club–neuroendocrine axis, underscoring the need for deeper sequencing or targeted enrichment in future studies. Second, while our IL-13–based in vitro model recapitulates key features of epithelial remodeling, it do not fully capture the complex *in vivo* interactions between the immune system and the epithelium. Incorporating immune components or utilizing airway-on-chip systems may enhance physiological relevance.

In conclusion, this work establishes an hPSC-derived club cell-enriched airway epithelial sheet platform that enables detailed analysis of epithelial identity, lineage plasticity, and disease-associated remodeling. By delineating conserved and previously unrecognized differentiation trajectories and providing a tractable system for therapeutic modeling, our study advances both fundamental understanding of human airway epithelial biology and the translational potential of hPSC-derived airway models.

## Acknowledgments

We thank S. Matsuo, Y. Maeda, S. Takahashi, and K. Ushitora for assistance with culture experiments, including routine medium changes. We are grateful to K. Okamoto-Furuta, H. Kohda, and T. Katsuno (Division of Electron Microscopic Study, Center for Anatomical Studies, Kyoto University) for their expert support with electron microscopy. We also thank M. Narita for assistance with secretome analysis and K. Deguchi, and S. Sakurai for their valuable assistance with bulk RNA-seq and single-cell RNA-seq analyses. We thank to Single-Cell Genome Information Analysis Core (SignAC) in ASHBi for sequencing.

## Author contributions

N.S., S.K, and S.G. conceived and designed the study. N.S., N.F., K.A., M.T., T.T., K.K., M.I., T.Y. and S.G. performed the experiments. N.S., N.F., S.K, M.I., T.Y. and S.G. analyzed the data. N.S., N.F., S.K, and S.G. wrote the manuscript.

## Figures

**Extended Data Movie 1:** Bright-field movie of a multiciliated epithelial cell sheet derived from GFP-positive cells in SCGB1A1 reporter hPSCs, analyzed by high-speed video microscopy (HSVM).

**Extended Data Table 1:** Contents of each iPSC differentiation medium.

|  | Endodermization<br>medium (Step1) | Anteriorization<br>medium (Step2) | Ventralization<br>medium (Step3) | Amplified<br>medium (Step4) |
| --- | --- | --- | --- | --- |
| Basal<br>Medium | RPMI<br>B27 supplement (2 %)<br>Penicillin/Streptomycin<br>(50 U/ml) | DMEM / F12<br>Glutamax<br>B27 supplement (2 %)<br>L-ascorbic acid (0.05 mg/ml)<br>monothioglycerol (0.4 mM)<br>Penicillin/Streptomycin (50 U/ml) |  |  |
| Chemicals<br>and<br>Cytokines | Activin A (100 ng/ml)<br>CHIR99021 (1.0 μM)<br>Y-27632 (Day0; 10 μM)<br>*sodium butyrate (NaB)*<br>(Day 1, 2 and 4) | Noggin (100 ng/ml)<br>SB431542 (10 μM) | ATRA (depends on cell lines)<br>B2-3 iPSCs: 0.05 μM,<br>SCGB1A1 reporter iPSCs: 0.1 μM,<br>585A1 iPSCs and 604A1 iPSCs: 0.5 μM<br>BMP4 (20 ng/ml)<br>CHIR99021 (3.0 μM) | CHIR99021 (3.0 μM)<br>FGF10 (10 ng/ml)<br>KGF (10 ng/mL)<br>DAPT (20 μM) |
|  | PAEPC medium (Step5) |  | PAEC medium (Step6-3D and 3D-ALI) |  |
| Basal<br>Medium | DMEM / F12<br>Glutamax<br>B27 supplement (2 %)<br>L-ascorbic acid (0.05 mg/ml)<br>monothioglycerol (0.4 mM)<br>Penicillin/Streptomycin (50 U/ml) |  | PneumaCult-ALI Basal Medium<br>PneumaCult-ALI Supplement<br>PneumaCult-ALI Maintenance Supplement |  |
| Chemicals<br>and<br>Cytokines | CHIR99021 (3.0 μM)<br>FGF10 (100 ng/ml)<br>Y-27632 (10 μM) | | Heparin (4 μg/ml)<br>Hydrocortisone (1.0 μM)<br>DAPT (10 μM)<br>Y-27632 (10 μM) | |

**Extended Data Table 2:** Antibodies used in this present study.

| Species | Antibodies | Company | Catalog No. | Dilution |
| --- | --- | --- | --- | --- |
| For isolating NKX2-1<sup>+</sup> cells |  |  |  |  |
| Rat | Anti-CPM | In house | N/A | 1:200- |
| - | Anti-rat IgG-MicroBeads | Miltenyi Biotec | 130-047-401 | 1:5- |
| For immunofluorescence staining (Primary Antibodies) |  |  |  |  |
| Rat | Anti-SCGB1A1 (CCSP) | R&D systems | MAB4218 | 1:25 |
| Chicken | Anti-Green Fluorescent Protein (GFP) | Aves labs | GFP-1020 | 1:500 |
| Rabbit | Anti-SCGB3A2 (UGRP) | Kindly provided by Prof. Dr kimura | N/A | 1:2000 |
| Mouse | Anti-EPCAM | Santa Cruz Biotechnology | sc-66020 | 1:500 |
| Mouse | Anti-Acetylated tubulin | Sigma-Aldrich | T7451 | 1:4000 |
| Mouse | Anti-FOXJ1 | eBioscience | 14-9965-82 | 1:500 |
| Mouse | Anti-MUC5AC | Thermo Scientific | MS-145-P | 1:100 |
| Mouse | Anti-MUC5B | Santa Cruz Biotechnology | sc-21768 | 1:25 |
| Goat | Anti-SYP | Santa Cruz Biotechnology | SC-7569 | 1:100 |
| Rabbit | Anti-SPC | Santa Cruz Biotechnology | sc-13979 | 1:100 |
| Mouse | Anti- SPB | Thermo Scientific | MS-704-P0 | 1:500 |
| Rabbit | Anti- proSPB | Millpore | AB3432 | 1:1000 |
| Rat | Anti- Ki67 | Thermo Scientific | 14-5698-82 | 1:500 |
| Rabbit | Anti- CCNO | Thermo Scientific | PA5-148399 | 1:200 |
| For immunofluorescence staining (Secondary Antibodies) |  |  |  |  |
| Donkey | anti-goat IgG-Alexa Fluor 488 | Thermo Fisher Scientific | A11055 | 1:500 |
| Goat | anti-mouse IgG-Alexa Fluor 488 | Thermo Fisher Scientific | A21202 | 1:500 |
| Goat | Anti-mouse IgG2b-Alexa Fluor 488 | Thermo Fisher Scientific | A21141 | 1:500 |
| Donkey | Anti-rabbit IgG-Cy3 | Jackson ImmunoResearch | 711-165-152 | 1:500 |
| Goat | Anti-mouse IgG1-Cy3 | Jackson ImmunoResearch | 115-165-205 | 1:500 |
| Donkey | Anti-rat IgG-Alexa Fluor 647 | Thermo Fisher Scientific | A48272 | 1:500 |

**Extended Data Table 3:**
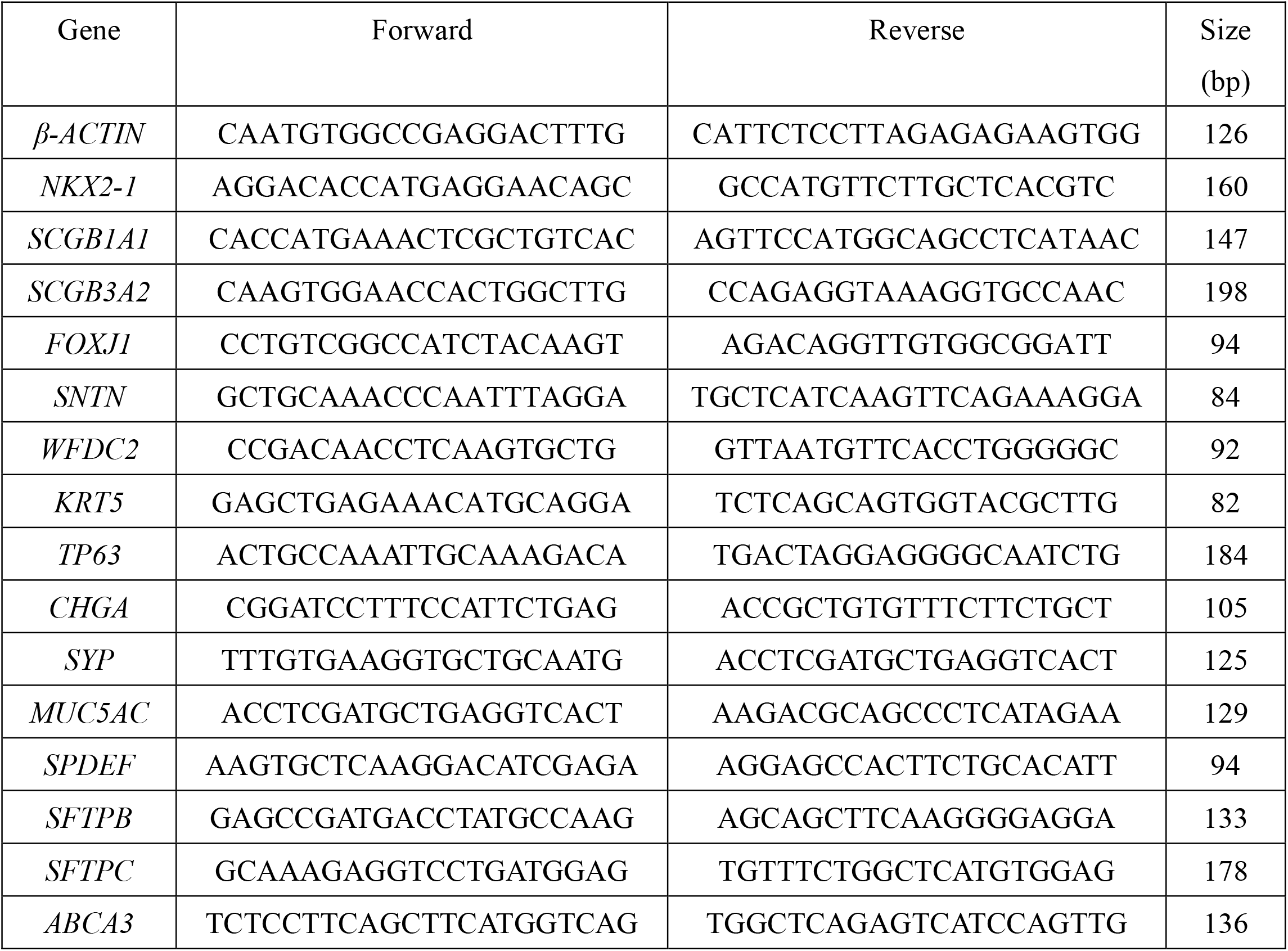
Primers for qRT-PCR used in this study.

---

**Extended Data Fig. 1:**
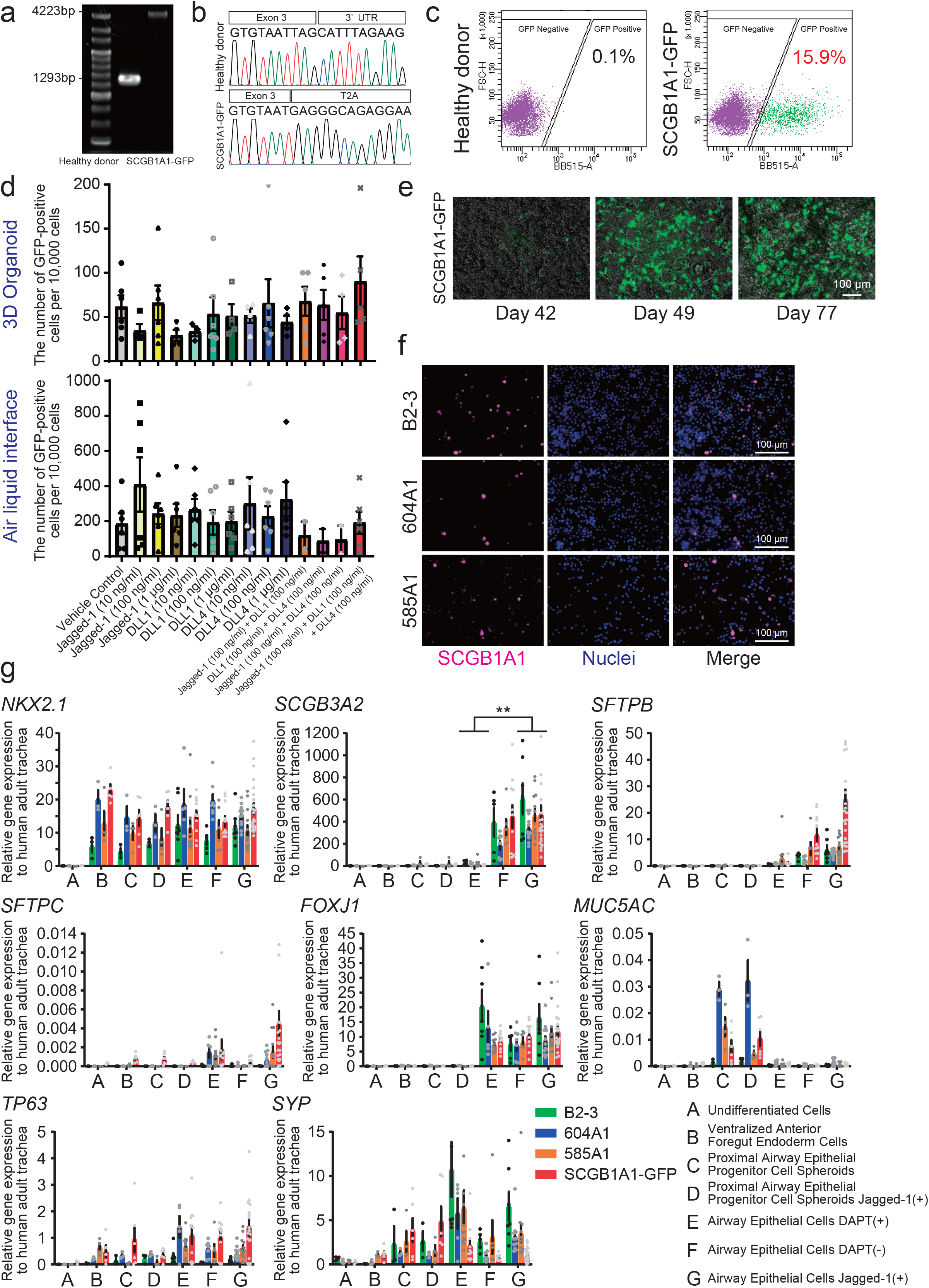
Supplementary characterization and validation of the differentiation protocol for SCGBlAl-positive club cells derived from hPSCs, related to Fig. 1. (a) Genomic validation of SCGB1A1 reporter knock-in by agarose gel electrophoresis. (b) Sanger sequencing confirmation of genomic integration of the SCGB1A1 reporter. (c) Representative flow cytometric profiles of GFP expression during club cell differentiation of SCGB1A1 reporter cells. (d) Comparative flow cytometric analysis of GFP expression under different Notch signaling activation conditions in 3D organoid and ALI cultures. (e) Timecourse analysis of GFP expression during club cell differentiation of the SCGB1A1 reporter cells (mean ± SEM, *n* ≥ 2 independent experiments). (f) Immunofluorescence validation of SCGBlAl-positive club cells in differentiated epithelial sheets derived from multiple hPSC lines. (g) RT-qPCR validation of additional cell markers of the club cell differentiation protocol across multiple hPSC lines (mean ± SEM, *n* ≥ 3 independent experiments; two-way ANOVA with Dunnett’s multiple comparisons test; \*\**p* < 0.01).

**Extended Data Fig. 2:**
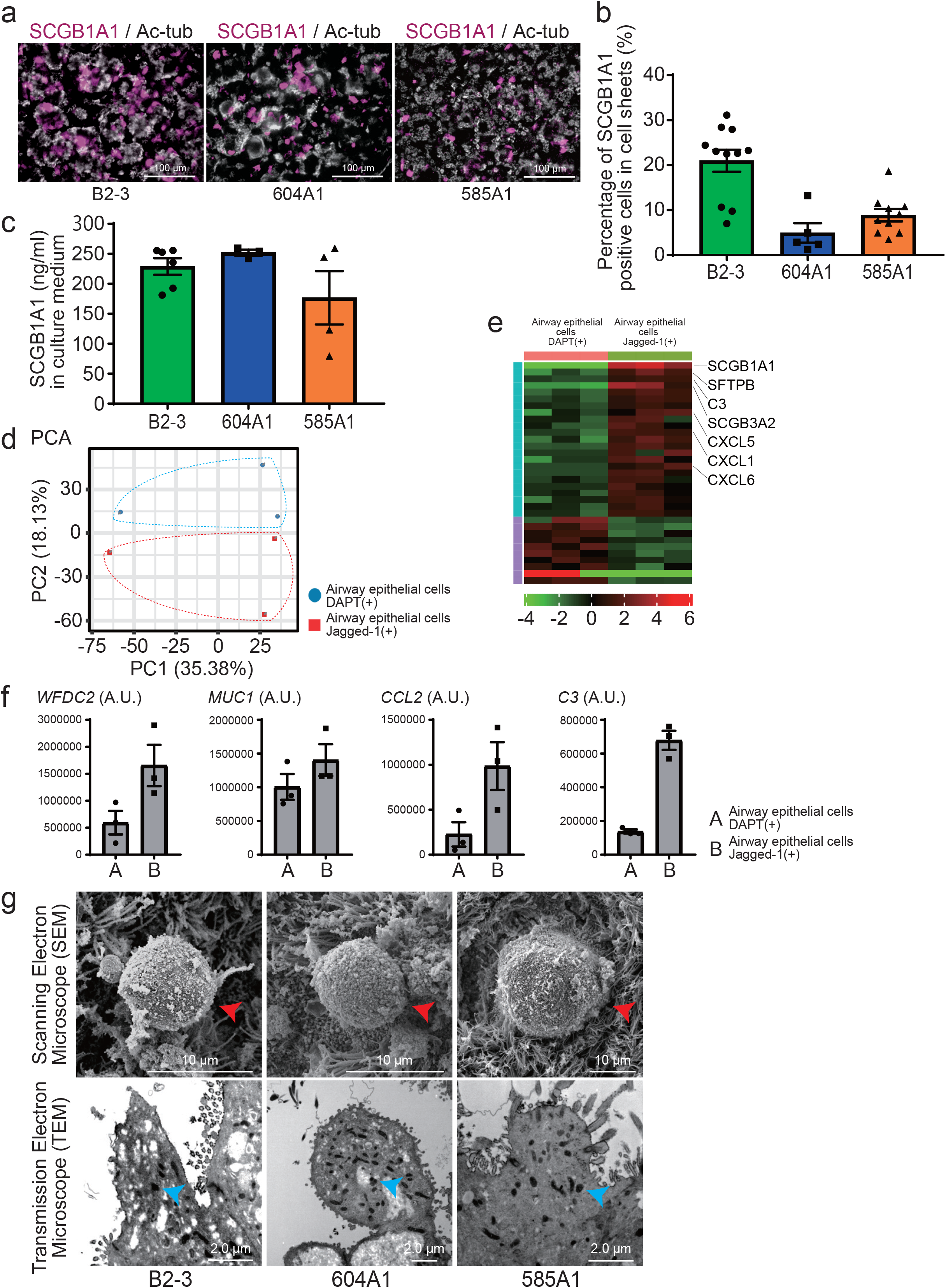
Supplementary validation of structural and functional maturation of the club cell-enriched airway epithelial sheet across multiple hPSC lines, related to Fig. 2. (a) Scattered distribution of SCGBlAl-positive club cells in club cell-enriched airway epithelial sheets derived from multiple hPSC lines. (b) Quantification of the proportion of scattered SCGBlAl-positive club cells in club cell-enriched airway epithelial sheets derived from multiple hPSC lines (mean ± SEM, *n* ≥ 3 independent experiments). (c) Evaluation of SCGB1A1 secretory function in club cell-enriched airway epithelial sheets derived from multiple hPSC lines by ELISA (mean ± SEM, *n* ≥ 3 independent experiments). (d) Principal component analysis of secretome profiles comparing the club cell-enriched airway epithelial sheet with Jagged-1 supplementation to that with Notch inhibitor, DAPT supplementation. (e) Heatmap visualization of secretome profiles comparing the club cell-enriched airway epithelial sheet and the airway epithelial sheet with DAPT supplementation. (f) Supplementary profiling of secreted proteins from the club cell-enriched airway epithelial sheet and airway epithelial sheet with DAPT supplementation (mean ± SEM, *n* = 3 independent experiments). Values are presented in protein area (A.U.), derived from mass spectrometry-based proteomics. (g) Ultrastructural characterization of the club cell-enriched airway epithelial sheets derived from multiple hPSC lines by transmission and scanning electron microscopy.

**Extended Data Fig. 3:**
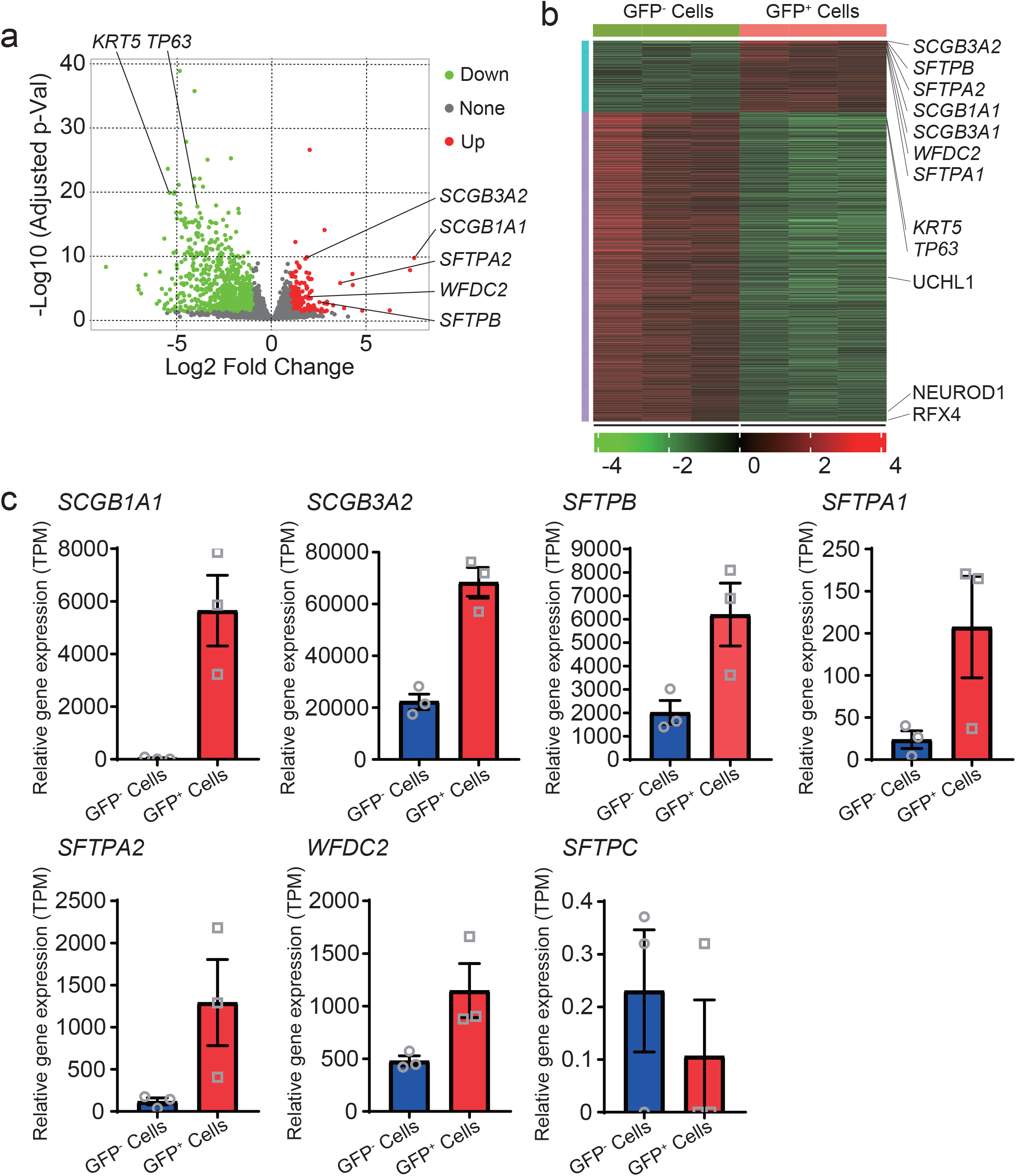
Supplementary RNA-seq analyses and additional validation of the bipotency of club cells derived from the club cell-enriched airway epithelial sheet, related to Fig. 2. (a) Volcano plot of RNA-seq data from GFP-positive and GFP-negative populations. (b) Heatmap visualization of RNA-seq data from GFP-positive and GFP-negative populations. (c) Expression profiles of club cell -associated genes derived from RNA-seq analysis of GFP-positive and GFP-negative populations (mean ± SEM, *n* = 3 independent experiments)

**Extended Data Fig. 4:**
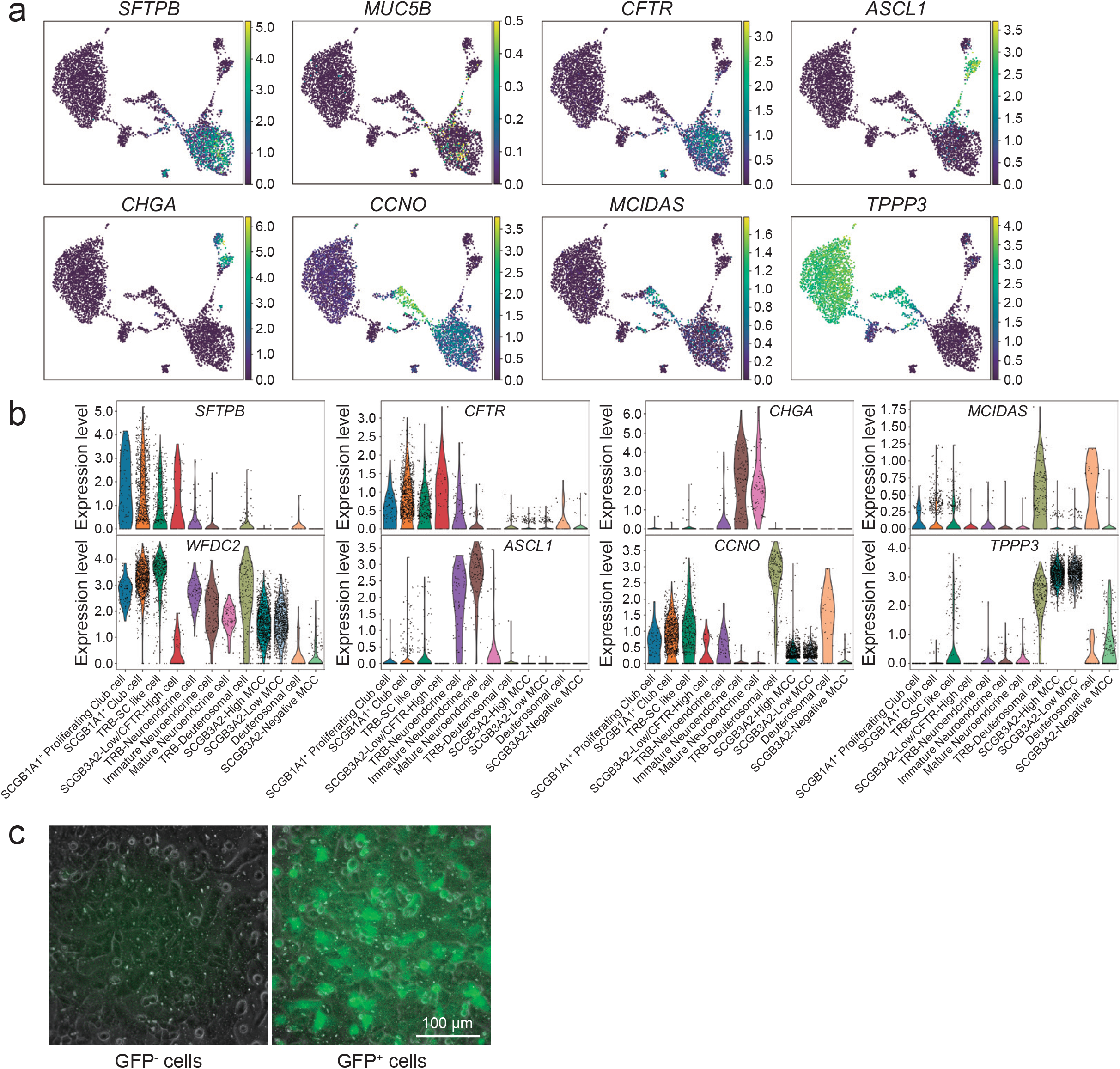
scRNA-seq analyses and immunofluorescence validation of SCGB3A2-positive club cells, TRB-neuroendocrine cells, and multiciliated epithelial cells identified by single-cell transcriptomic analysis, related to Fig. 3 and Fig. 4. (a) Feature plot analysis of additional marker genes for club cells, proliferative cells, deuterosomal cells, multiciliated epithelial cells, and neuroendocrine cells, not shown in Fig. 4c. (b) Violin plot analysis of additional marker gene expression across cell clusters identified in the club cell-enriched airway epithelial sheet, complementing Fig. 4e. (c) Validation of GFP expression in FACS-isolated GFP-positive and GFP-negative populations.

**Extended Data Fig. 5:**
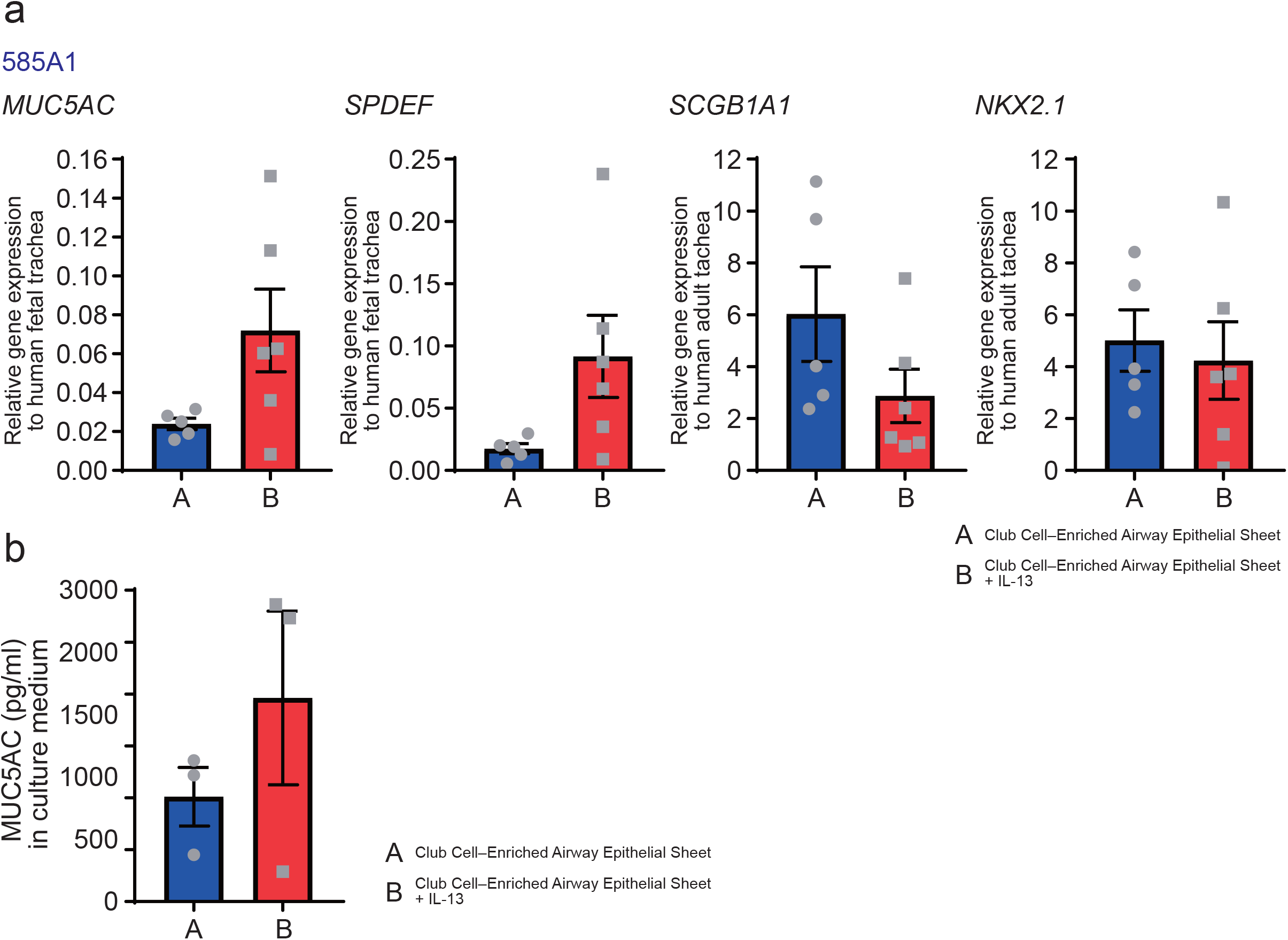
Independent validation of IL-13 -induced epithelial remodeling in airway epithelial sheets, related to Fig. 5. (a) RT -qPCR analysis of goblet and club cell marker gene expression in control and IL-13 -treated airway epithelial sheets from additional hPSC lines (mean ± SEM, n ≥ 3 independent experiments). (b) ELISA quantification of MUC5AC secretion in control and IL-13 -treated airway epithelial sheets from additional hPSC lines (mean ± SEM, n ≥ 2 independent experiments).

